# Targeting Protease-activated Receptor 4 (PAR4) Protects Against Acute Kidney Injury (AKI) in Ischemia–Reperfusion Injury

**DOI:** 10.64898/2026.03.27.714572

**Authors:** Emma Webb, Shirong Cao, Yu Pan, MingZhi Zhang, Raymond Harris, Olivier Boutaud, Jacob L. Bouchard, Carrie K. Jones, Craig W. Lindsley, Heidi E. Hamm

## Abstract

Acute kidney injury (AKI) is a serious and common clinical syndrome that currently has no effective treatment. Emerging evidence links coagulation pathways to kidney injury, particularly through coagulation proteases. Protease-activated receptors (PARs) are a family of G-protein coupled receptors (GPCRs) that are activated by proteolytic cleavage of their N termini, exposing a tethered ligand that initiates receptor signaling. PARs have been shown to play a major role in inflammation, vascular regulation, and tissue injury. PARs play key roles in inflammation, vascular regulation, and tissue injury. Previous work from the Hamm laboratory demonstrated that PAR4 contributes to AKI progression, as PAR4 knockout mice were protected in both unilateral ureteral obstruction and ischemia-reperfusion–based models of kidney disease.

In this study, we investigated the potential of a PAR4 antagonist, VU6073819, at mitigating AKI progression in an ischemia-reperfusion injury (IRI) mouse model. PAR4 antagonism not only alleviated kidney injury and inflammatory response, but it significantly improved the survival. These findings identify PAR4 as a promising therapeutic target for AKI.

## Introduction

Acute kidney injury (AKI) is a common clinical syndrome characterized by a rapid decline in renal function. AKI occurs in approximately 5–7% of hospitalized patients and in up to 50–60% of critically ill individuals^1^. The condition arises from diverse etiologies including impaired renal perfusion, direct tissue injury, and obstruction of urinary outflow. AKI is associated with prolonged hospitalization, increased healthcare costs, and elevated mortality, and it significantly increases the risk of cardiovascular complications and progression to chronic kidney disease^1^. Despite its clinical burden, there are currently no targeted therapies for AKI, and treatment remains largely supportive.

Emerging evidence suggests that coagulation proteases contribute to kidney injury beyond their traditional roles in hemostasis and thrombosis^2^. Protease-activated receptors (PARs) are a family of G-protein– coupled receptors that are activated through proteolytic cleavage of their extracellular N-terminus, exposing a tethered ligand that initiates receptor signaling^3,4^. This unique activation mechanism links protease activity to cellular responses in inflammation, vascular regulation, and tissue injury. Several PAR family members are expressed in renal cell types, suggesting that protease signaling may influence renal physiology and pathology.

Among these receptors, protease-activated receptor 4 (PAR4) has emerged as a potential mediator of inflammatory injury^5–7^. PAR4 is highly expressed on platelets but has also been detected in endothelial and immune cells, where it contributes to pro-inflammatory signaling and leukocyte recruitment^8–10^. Recent work from the Hamm and Harris laboratories demonstrated that PAR4 expression is markedly induced in injured kidneys in mouse models of unilateral ureteral obstruction and ischemia-reperfusion injury^11^. In these studies, PAR4 localized to distal tubular epithelial cells (**Figure 1**) and knocking out PAR4 protected mice from renal fibrosis, inflammation, and functional decline^11^. These findings suggest that PAR4 signaling contributes to kidney injury and may represent a therapeutic target in AKI.

**Figure 1.**
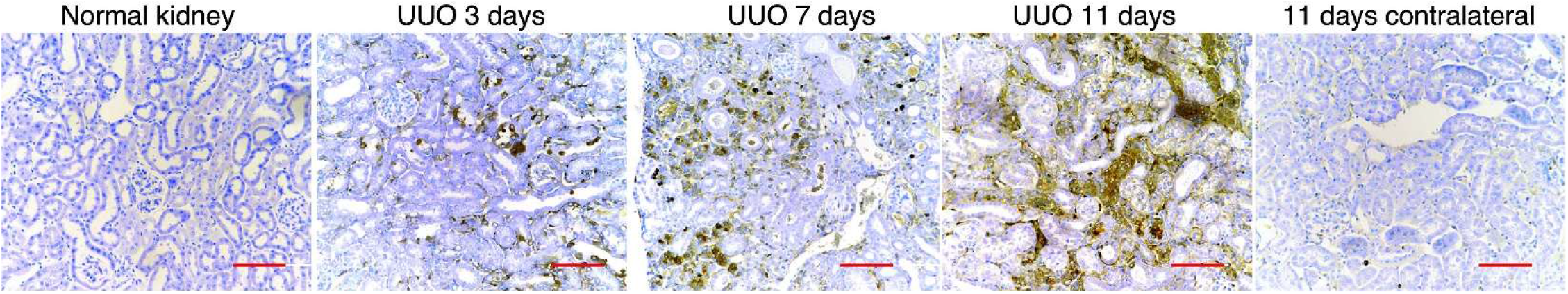
PAR4 expression increases in kidney distal tubule epithelial cells post unilateral ureteral obstruction (UUO) surgery. Representative images of PAR4 induction following UUO in WT C57/Bl6 mice over time (*n* = 10 in wild-type mice). PAR4 staining was minimal in control kidney and in the contralateral control kidney remained low even at 11 days. Scale bar = 50 μm. UUO, unilateral ureter obstruction; WT, wild type. *n* indicates the number of mice for each experiment.

In the present study, we tested whether pharmacological antagonism of PAR4 protects against AKI. Using a mouse model of ischemia–reperfusion injury, we evaluated the effects of the selective PAR4 antagonist, VU6073819, on renal function, inflammatory responses, and survival. Our findings demonstrate that PAR4 antagonism attenuates kidney injury and improves survival, supporting PAR4 as a promising therapeutic target for AKI.

## Methods

All studies were conducted in adult male C57BL/6J mice. Animals were under a 12/12 h light-dark cycle with water available ad libitum. All animal experiments were approved by the Vanderbilt University Animal Care and Use Committee, and experimental procedures conformed to guidelines established by the National Institutes of Health Guide for the Care and Use of Laboratory Animals. All efforts were made to minimize animal suffering and the number of animals used.

### IRI Model

8- to 12-week-old C57BL mice were anesthetized with 2% isoflurane, and their left kidneys are exposed via a left flank incision. Renal pedicle clamping occurs for 30 minutes, inducing the ischemia-reperfusion injury (IRI). 12 hours prior of injury and every day after, mice were administered either the vehicle or a PAR4 antagonist at 1 mg/kg and 5 mg/kg BID via IP administration. Blood samples were collected on days 2, 3, and 6. A subset of animals was sacrificed at day 3 to assess early injury markers, while the remaining mice were sacrificed at day 8. Mice were dosed BID with vehicle (10% Tween80:sterile water) or with VU6073819 at 1 mg/kg or 5 mg/kg. Mice were randomized to treatment groups and investigators were blinded during analysis. Blood samples were taken on day 2, 3, and 6 of the study to measure BUN and drug concentration. Mice were sacrificed 8 days following IRI for histological examination of fibrosis and RT-PCR analysis of genes related to fibrosis and inflammation.

### Quantitative immunofluorescence/immunohistochemistry staining

Kidney tissue was immersed in fixative containing 3.7% formaldehyde, 10 mM sodium m-periodate, 40 mM phosphate buffer, and 1% acetic acid. The tissue was dehydrated through a graded series of ethanols, embedded in paraffin, sectioned (5 μm), and mounted on glass slides. Rat anti-F4/80 (BioRad, MCA497) and Rabbit anti-Ly6g (1A8, Abcam, ab238132) were used for immunostaining. Antigen retrieval in the deparaffinized sections was performed with citrate buffer by microwave heat for 10 min and the slides were then blocked with 10% normal donkey serum for 1 hour at RT followed by incubation with primary antibodies overnight at 4°C. For immunofluorescence staining, the sections were incubated in two rounds of staining overnight at 4°C. Anti-rabbit or mouse IgG-HRP were used as secondary antibodies (Cell Signaling Technology). Each round was followed by tyramide signal amplification with the appropriate fluorophore (Alexa Flour 488 tyramide, Alexa Flour 647 tyramide or Alexa Flour 555 tyramide, Tyramide SuperBoost Kit with Alexa Fluor Tyramides, Invitrogen) according to its manufacturer’s protocols. PAR4 monoclonal Ab was used as previous report^11^. DBA was used as collecting duct marker, LTL as proximal tubule marker and DAPI as a nuclear stain. Sections were viewed and imaged with a Nikon TE300 fluorescence microscope and spot-cam digital camera (Diagnostic Instruments), followed by quantification of cells/field using Image J software (NIH, Bethesda, MD). IOD were calculated in more than 30 fields per mouse or 10 fields per cell slide and expressed as arbitrary units or percentage per field by two independent investigators.

### RT-PCR

Total RNAs from kidneys were isolated using TRIzol® reagent (Invitrogen). SuperScript IV First-Strand Synthesis System kit (Invitrogen) was used to synthesize cDNA from equal amounts of total RNA from each sample. Quantitative RT-PCR was performed using TaqMan real-time PCR (7900HT, Applied Biosystems). The Master Mix and all gene probes were also purchased from Applied Biosystems. The probes used in the experiments included mouse GAPDH (Mm99999915), collagen I (*Col1a1*, Mm00801666), collagen III (*Col3a1*, Mm01254476), collagen IV (*Col4a1*, Mm01210125), fibronectin 1 (*Fn1*, Mm01256744), Vimentin (*Vim*, Mm01333430), CTGF (*Ctgf*, Mm01192933), TGF-β1 (Tgfb1,Mm00441726), F4/80 (*Emr1*, Mm00802529), *Cd68* (Mm03047343), *Ly6g* (Mm04934123), *Havcr1* (*Kim-1*, Mm00506686), *Lcn2(Nagl*, Mm01324470*), Tnf* (Mm99999068), *Il1a (*Mm00439620), *Il1b* (Mm00434228), *Ccl2* (Mm00441242), *Ccl3* (Mm00441258), and *Il23a* (Mm00518984).

### Histology analysis

Periodic acid Schiff (PAS) stained slides were evaluated for tubular injury without knowledge of the identity of the various groups. A semiquantitative evaluation was used to evaluate the degree of tubular injury. Each low power field of tubules on a single section was graded from 0–5, with 1, 2, 3, 4 and 5 representing tubular injury, <10, 10-25, 25–50, 50–75, or >75% of the injury area, respectively. Tubular injury includes dilation of the tubules and flattening of the tubular epithelium, tubular casts, fragments of cells or necrotic epithelium in the tubular lumen, loss of brush border, loss of nuclei, and denudation of the basement membrane.

*Picrosirius red stain* was performed according to the protocol provided by the manufacturer (Sigma, St. Louis, MO, USA).

### Measurement of BUN

BUN was measured using a Urea Assay Kit (BioAssay Systems, Hayward, CA).

### Drug Quantification in Plasma

Concentrations of VU6073819 in plasma were quantified by liquid chromatography tandem mass spectrometry (LC-MS/MS). Plasma samples were centrifuged at 3,500 g for 5 min. A standard curve was generated by diluting the analyte DMSO stocks with blank plasma to a final concentration of 10,000 ng/mL, then serially diluting to 0.5 ng/mL. Quality controls were generated by a serial dilution of the 5,000 ng/mL standard curve solution in blank plasma to obtain 3 concentrations of 500, 50, and 5 ng/ml. 20 uL of plasma, blank plasma, standard curve, and QC samples were loaded in a V-bottom 96-well plate. 120 uL of acetonitrile containing 0.05 uM carbamazepine (internal standard) was added to each well, and the plate was centrifuged at 3,500 g for 5 min. 60 uL of the supernatant of each well (protein-free) was transferred to a new 96-well plate containing 60 uL of water. The plates were sealed for LC-MS/MS analysis.

### Statistical Analysis

Statistical analyses were performed using GraphPad Prism (version 11.0.0; GraphPad Software, San Diego, CA, USA). Data are presented as mean ± standard error of the mean (SEM) unless otherwise indicated. Comparisons among multiple groups were performed using one-way or two-way analysis of variance (ANOVA) followed by appropriate post hoc multiple comparison tests (e.g., Tukey’s or Bonferroni’s test). Survival curves were analyzed using the Kaplan–Meier method with log-rank testing. A p value < 0.05 was considered statistically significant.

## Results

This study investigated the effect of PAR4 antagonism in a mouse model of AKI induced by IRI. Outcomes used to evaluate protection against AKI included kidney function, inflammatory markers, and overall survival. Mice treated with VU6073819 showed a dose-dependent response in protection against changes in body weight (**Figure 2A**), kidney weight, and survival (**Figure 2B**). Survival was significantly improved in the VU6073819-treated group. Six of ten vehicle-treated mice died prior to study termination, whereas all mice receiving 5 mg/kg VU6073819 survived to day 8.

**Figure 2.**
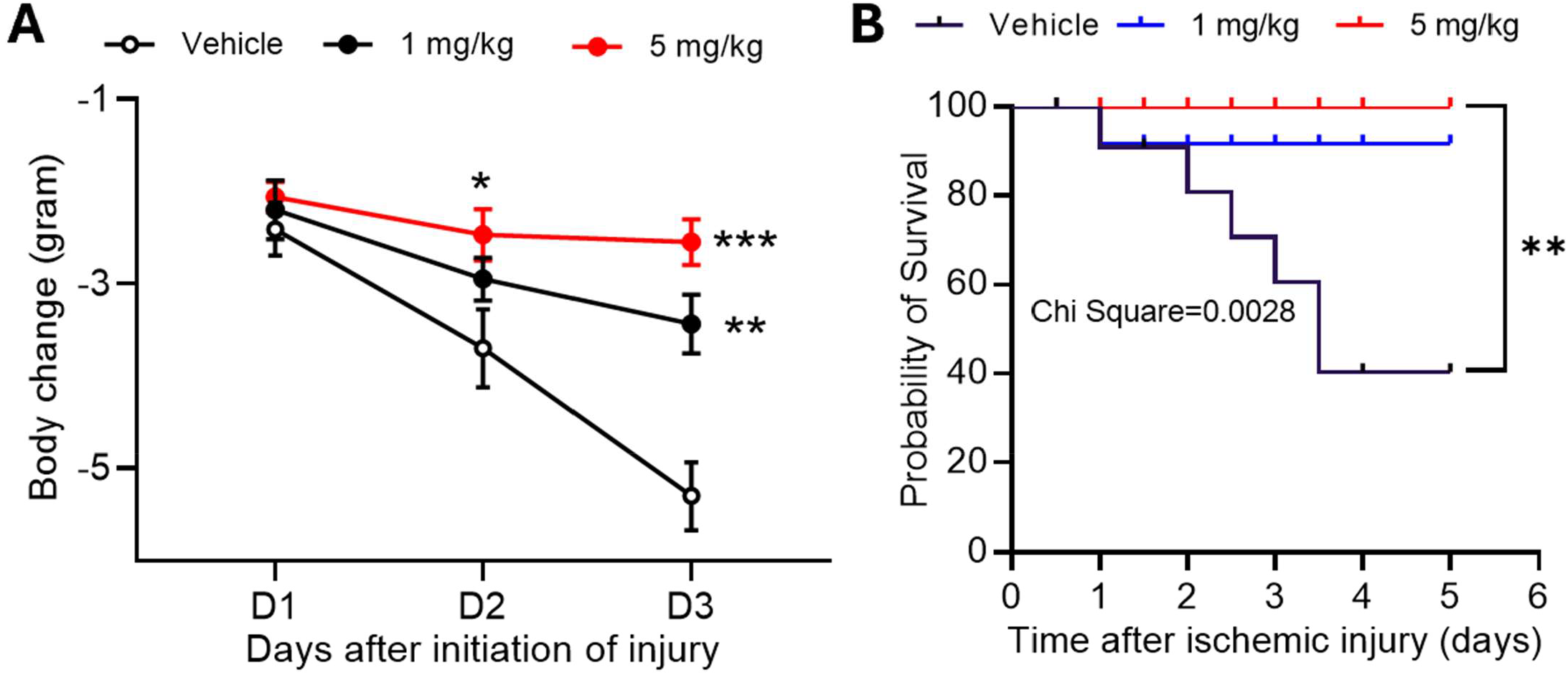
PAR4 antagonist protects WT mice against a mouse of AKI. WT mice were dosed with vehicle or PAR4 antagonist at 1mg/kg or 5 mg/kg by I.P. injection and BID. Blood samples were taken on day 2, 3, and 6 of the 8-day study. Mice were pre-treated with PAR4 antagonist or vehicle prior to ischemia-reperfusion injury (IRI). A) Dose dependent effect seen in body weight change and B) and survival curve analysis. Mean ± SEM, N = 10 per treatment group. Mean ± SEM; N = 10. Two-way ANOVA, Šídák’s post hoc shows significant difference in body weight change (A). Kaplan-Meier survival analysis, followed by Log-rank test shows significance between vehicle treated and 5 mg/kg treated mice in survival, providing a p-value. *(**P* ≤ 0.001).

VU6073819 treatment produced a dose-dependent improvement in renal function, reflected by reduced blood urea nitrogen (BUN) levels (**Figure 3**). This improvement was accompanied by decreased inflammatory and injury markers including Kim-1, Cd68, and Il1b (**Figure 4**). Plasma exposure at the end of the study was well above the IC_50_ needed to maintain therapeutic effect (**Figure 5**), indicating that at both 1 mg/kg and 5 mg/kg, there is sufficient drug exposure to elicit antagonism of PAR4, and a good PK/PD relationship.

**Figure 3.**
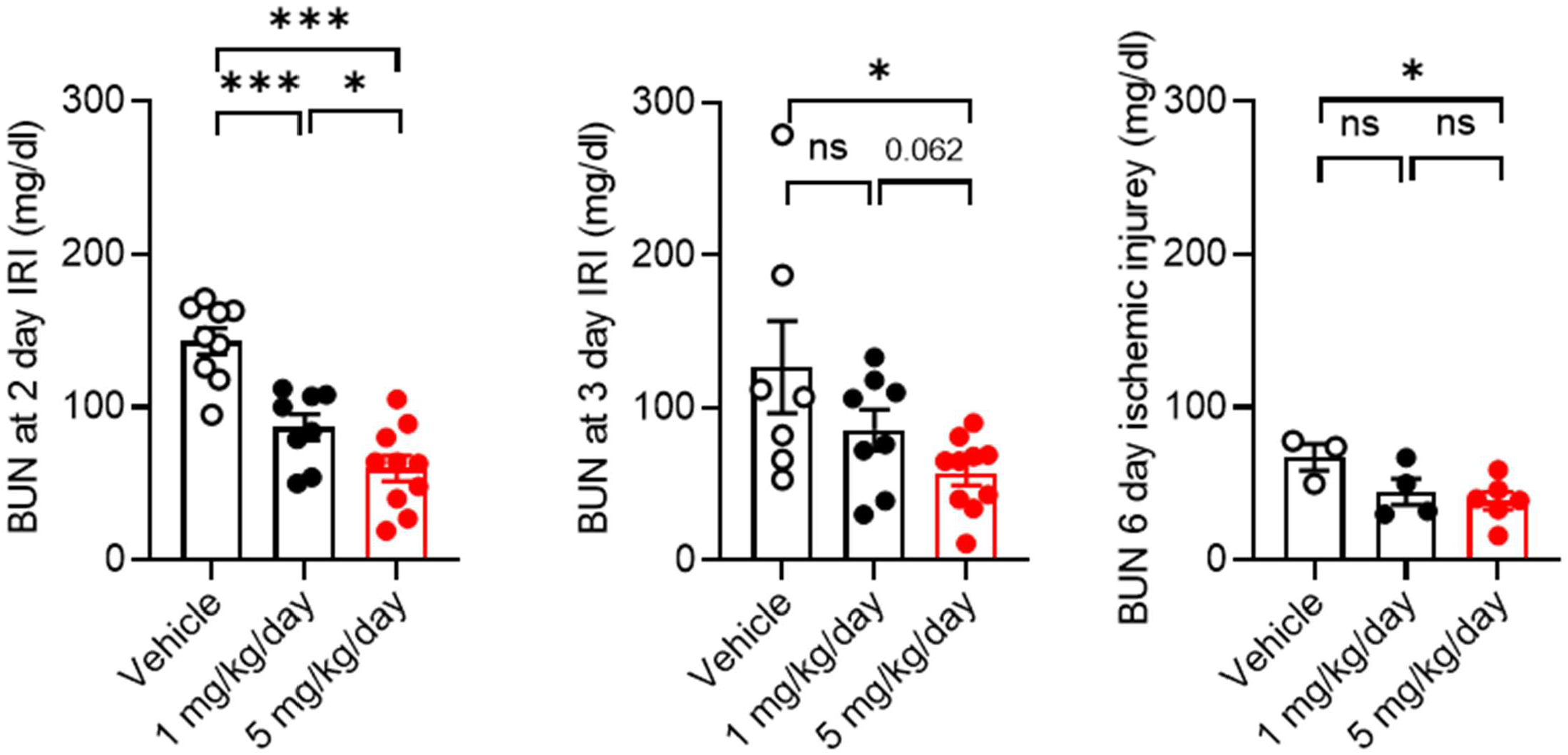
PAR4 antagonist, VU6073819, protects WT mice against kidney injury. Dose dependent effect in blood urea nitrogen (BUN) levels demonstrating protection of disease progression when administered VU6073819. Mean ± SEM, the number of animals/groups differed due to survival rate. 5 mg/kg treatment group was the only group to maintain all 10 mice from the start of the study. One-way ANOVA; Tukey post hoc analysis. \**P* ≤ 0.05, \*\*\**P* ≤ 0.001.

**Figure 4.**
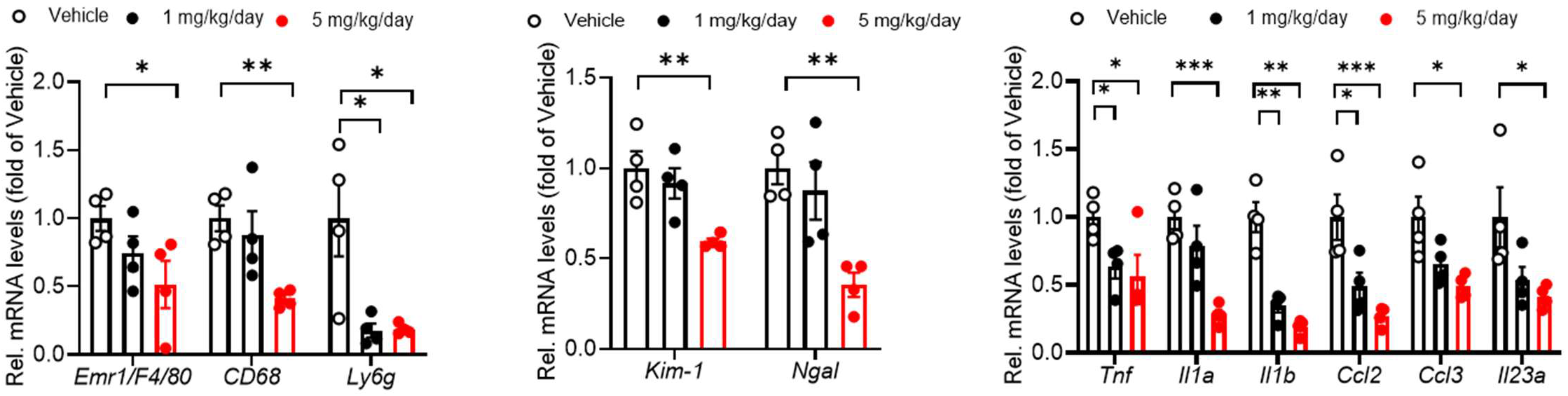
PAR4 antagonist, VU6073819, protects WT mice against kidney injury and inflammation markers. Dose dependent effect in immune cell markers, kidney injury markers, and inflammatory markers demonstrating protection of disease progression when administered VU6073819. Mean ± SEM, the number of animals/groups differed due to survival rate. 5 mg/kg treatment group was the only group to maintain all 10 mice from the start of the study. One-way ANOVA; Tukey post hoc analysis. \**P* ≤ 0.05, \*\**P* ≤ 0.01, \*\*\**P* ≤ 0.001.

**Figure 5.**
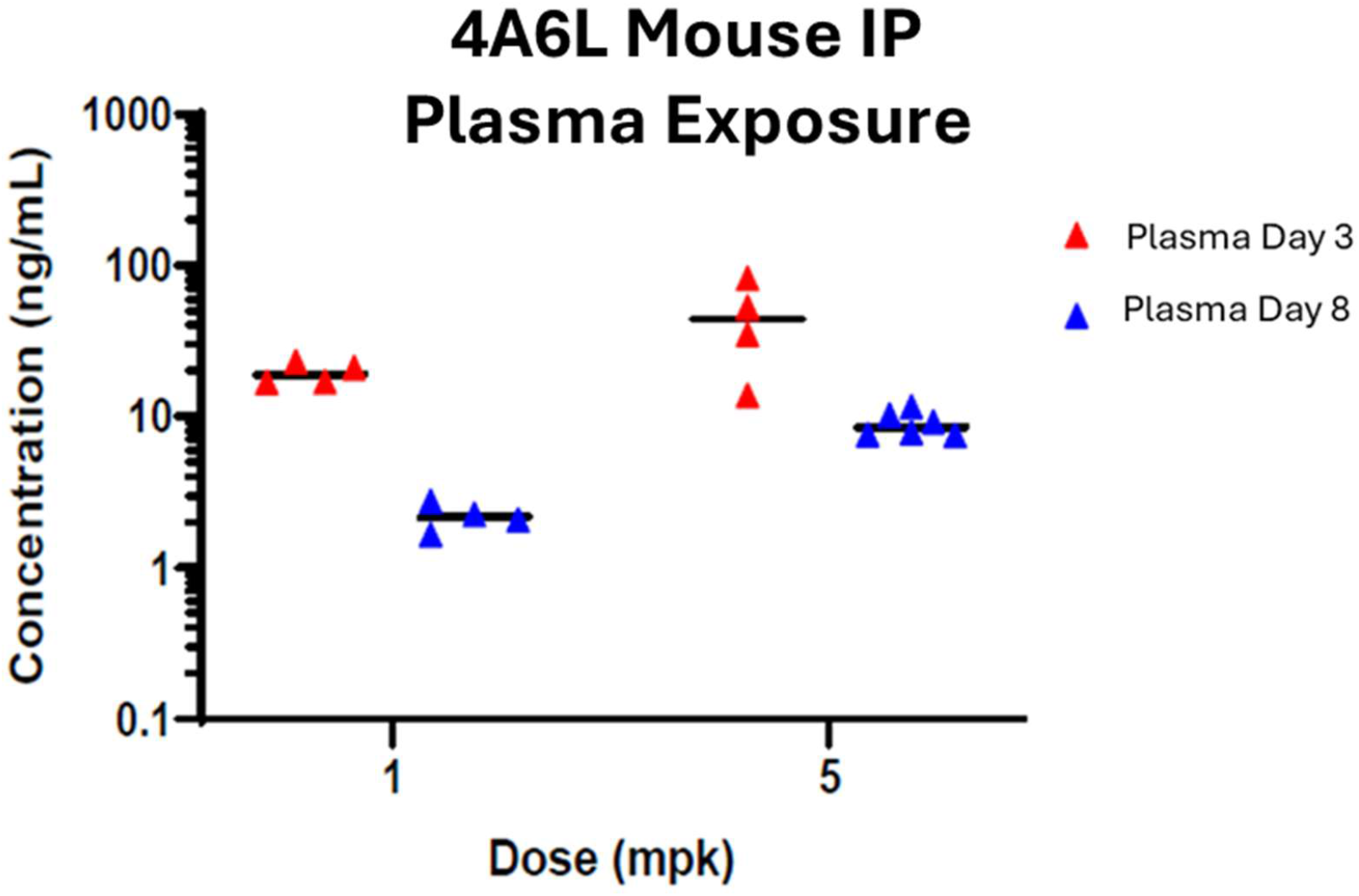
Plasma concentration of 4A6L for 1 mg/kg and 5 mg/kg doses at time of sacrifice, Day 3 and Day 8. The concentration of 4A6L was above the IC_50_ of the compound (IC_50_ = 45 nM). Mean ± SEM, N=8 in 1 mg/kg treatment group, and N=10 in 5 mg/kg treatment group.

Further analysis was conducted on the 5 mg/kg VU6073819 treated mice at the day8 time point. VU6073819 treated mice were protected against kidney tubule injury, indicated by periodic acid Schiff staining and blinded histological analysis (**Figure 6**).

**Figure 6.**
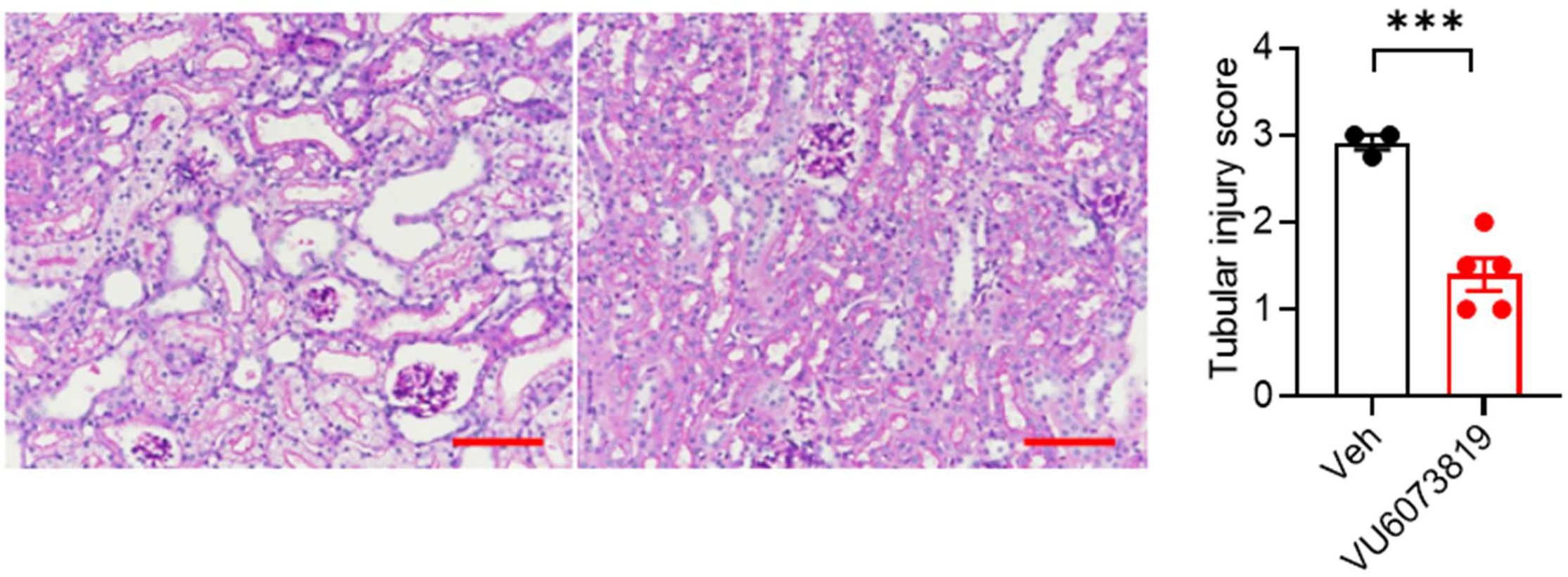
Kidney tubular injury score for 5 mg/kg VU6073819-treated mice at time of sacrifice, Day 8. Observed by Periodic acid Schiff staining and blinded histological analysis. Scale bar: 50 M. Mean ± SEM, N=5 in 5 mg/kg treatment group, and N=3 in vehicle treatment group. Welche’s t test; two-tailed; \*\*\**P* ≤ 0.001.

VU6073819 treatment also decreased neutrophil and macrophage infiltration to the kidney after injury compared to vehicle control (**Figure 7A** and **Figure 7B**, respectively). Both neutrophils and macrophages increase inflammatory response and can cause further damage at the site of injury^12–14^. By antagonizing PAR4, we see a reduction in inflammatory response, mitigating the damage done to the kidney.

**Figure 7.**
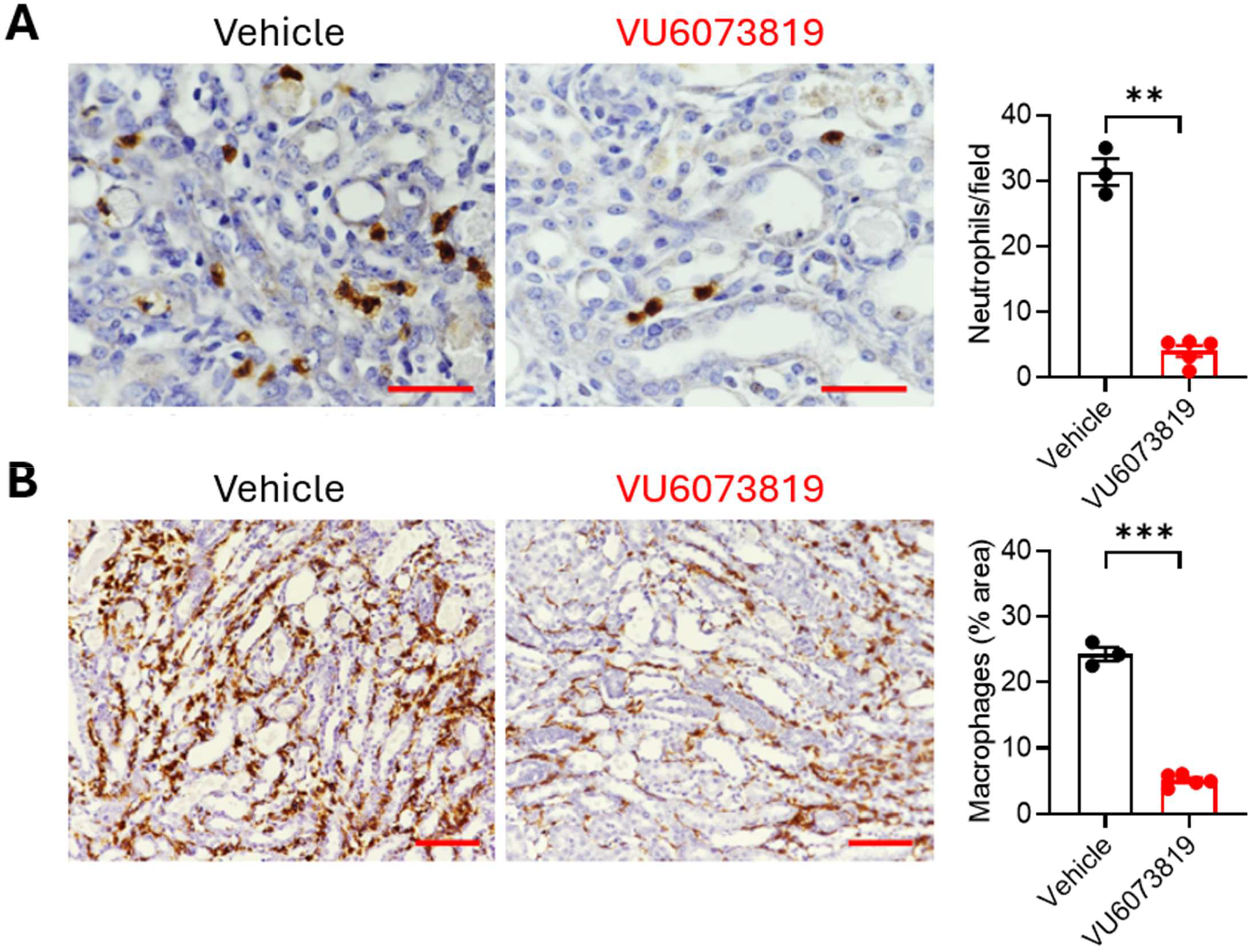
Immune cell infiltration quantification in kidneys after IRI. A) Neutrophil infiltration quantified by Ly6g staining of kidney sections and blinded histological analysis. Welche’s t test; two tailed; \*\**P≤0*.*01* B) Macrophage infiltration measured by F4/80 staining of kidney sections and blinded histological analysis. Welche’s t test; two tailed; \*\**P* ≤ 0.001.

Fibrosis was also reduced in VU6073819-treated mice compared to vehicle controls, as demonstrated by Picrosirius red staining (**Figure 8A**). This finding was supported by decreased expression of fibrosis-associated genes, including *VIM, FN1, CTGF*, and *TGFB1* (**Figure 8B**). Given that these genes are also involved in tissue repair, these results raise the possibility that PAR4 antagonism may influence normal repair processes.

**Figure 8.**
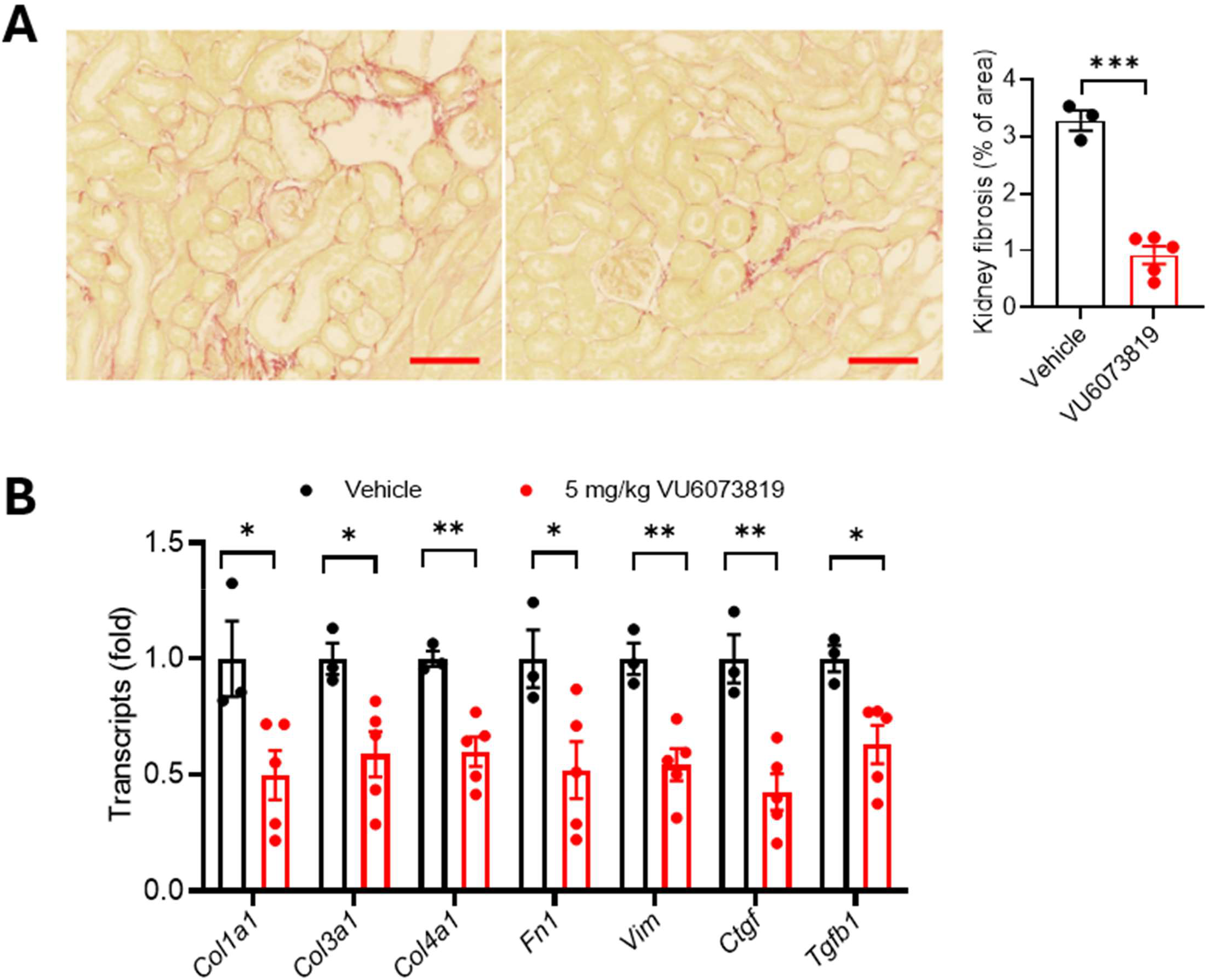
Fibrosis quantification of kidneys after IRI. A) Representative images for vehicle and VU6073819 fibrosis visualized by Sirius Red staining 8 days following IRI. Sirius Red quantification expressed as percent area shows decreased fibrosis in VU6073819 compared with vehicle mice. \*\*\**P* < 0.001 Welch’s *t* test, *n* = 3-5. Scale bar = 50 μm. *n* indicates the number of mice for each experiment. B) mRNA of fibrosis markers for vehicle and VU6073819 8 days following IRI. One-way ANOVA; Tukey post hoc analysis. *P* ≤ 0.05, **P ≤ 0.01, \*\*\**P* ≤ 0.001.

To confirm translational relevance, human kidneys were stained to look at PAR4 expression. In kidneys from patients with acute interstitial nephritis (AIN), PAR4 expression colocalized with the distal tubule marker DBA, indicating localization to the distal convoluted tubule (**Figure 9**). In contrast, minimal PAR4 expression was observed in normal human kidney tissue. These findings suggest that PAR4 expression is induced in human kidney disease and may contribute to tubular injury and inflammation. Inflammation in the distal tubule results in increased inflammatory cytokines, reactive oxygen species (ROS), and cell adhesion molecules, leading to tissue damage, apoptosis, and kidney atrophy. PAR4 expression in the distal tubule may be driving epithelial barrier breakdown and immune cell infiltration^15–18^.

**Figure 9.**
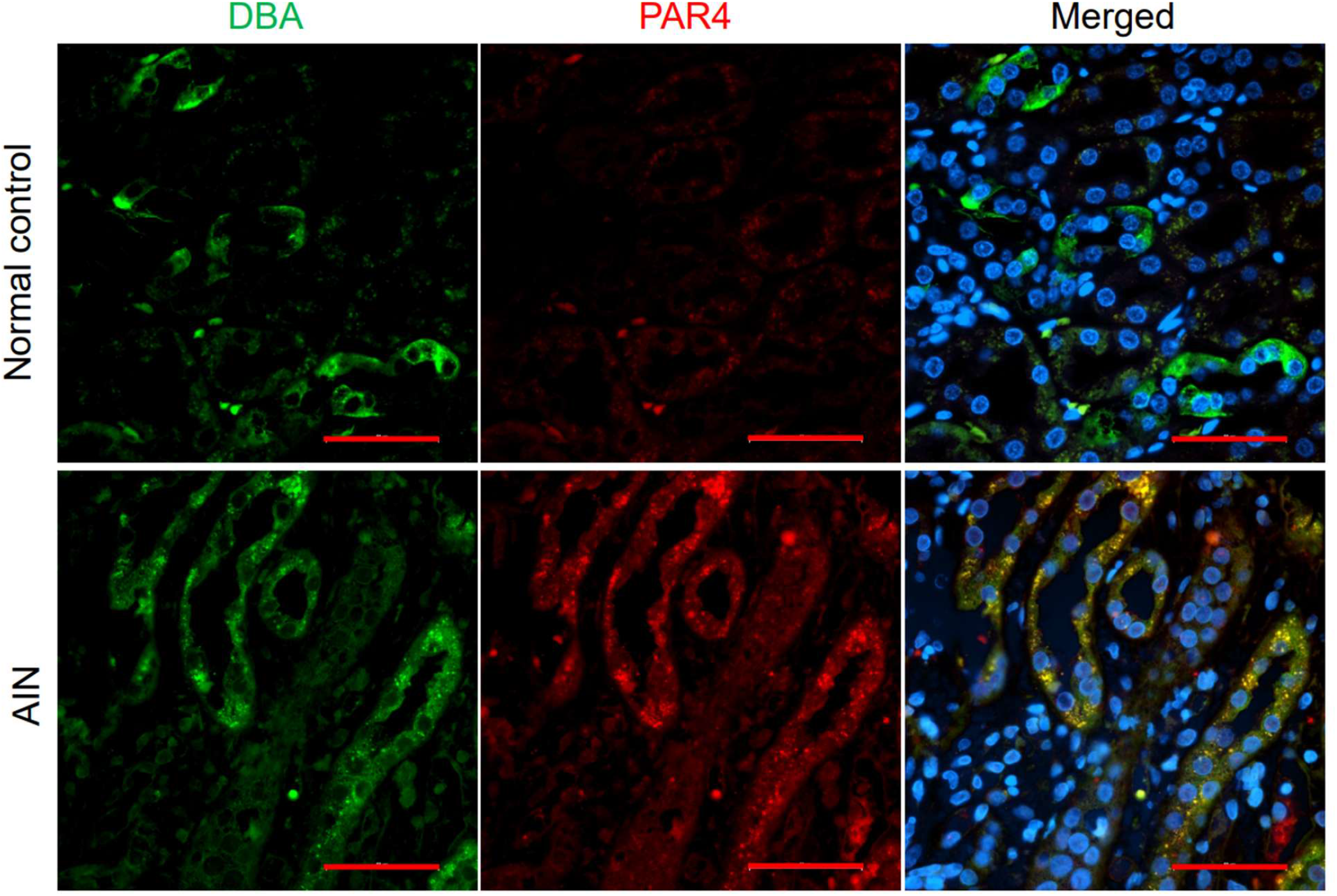
PAR4 expression localized to the renal collecting duct in human acute interstitial nephritis (AIN). Representative human kidney images stained with 14H6 PAR4 antibody, DBA collecting duct marker, and nucleus marker, DAPI. Scale bar: 50 M.

## Discussion

The PAR receptors have been investigated as potential players in renal diseases, with PAR2 and PAR3 being expressed in podocytes. PAR1 and PAR4 are mainly thought to play roles in thrombosis and hemostasis due to their high expression on platelets, however, they are also involved in pro-inflammatory signaling in many tissues. PAR4 expression has been reported on endothelial cells, leukocytes, microglia, and neurons. PAR4 knockout (PAR4 KO) studies have demonstrated a role of PAR4 in neutrophil homing and invasion of vascular injury, and PAR4-mediated neutrophil recruitment underlies tissue damage after ischemia-reperfusion injury in animal models of myocardial infarction and stroke, as well as inflammatory bowel disease, kidney disease, and arthritis^7,8,11,16,19–27^.

We previously showed an up-regulation of PAR4 in kidney convoluted distal tubule epithelial cells after unilateral ureteral obstruction, and PAR4 knockout protects against damage from UUO^11^. Here we show that inhibition of PAR4 activity protects against AKI, again showing clearly that PAR4 plays a role in AKI progression in the IRI mouse model of AKI. The PAR4 antagonist-treated mice were protected from disease progression and had overall better survival and better kidney function than vehicle control. There are still many questions about the biological role of PAR4 in AKI progression. PAR4 expression in convoluted distal tubule epithelial cells is almost undetectable; however, under disease conditions, PAR4 becomes expressed. The signaling mechanism involving PAR4 needs to be characterized to fully understand the pathological role of PAR4 in AKI. PAR4 is known to affect endothelial permeability and may be affecting epithelial cells in a similar manner, which would increase immune cell infiltration and perpetuate inflammation.

It is also unknown which protease activates PAR4 in this microenvironment. Thrombin has historically been considered the main activator of PAR4, but the kidney also expresses plasmin, kallikrein, trypsin-like proteases, cathepsin G, and MASP-1. All of these proteases have previously been shown to activate PAR4^9,28–34^. Tissue kallikrein was found to mediate pro-inflammatory pathway activation of PAR4 in proximal tubular epithelial cells in the kidney ^32^. Renal IRI leads to activation of the coagulation cascade, increase in infiltrating neutrophils, and secretion of proteases by damaged tubule epithelial cells. All of these processes involve proteases that can activate PAR4. Understanding how PAR4 is activated in the kidney will contribute to the understanding of PAR4’s role in AKI progression, as well as aid the development of more potent and specific PAR4 antagonists.

It would also be beneficial to understand which cell type expressing PAR4 is the major mediator of AKI progression. PAR4 is known to be expressed on platelets and endothelial cells, and we have shown PAR4 expression in the convoluted distal tubule epithelial cells in the UUO mouse model^11^. By treating the mouse systemically with a PAR4 antagonist, we are not able to distinguish if it’s the PAR4 on platelets, kidney epithelium, or vascular endothelium that is driving AKI progression in the IRI mouse model. The Hamm laboratory plans to utilize tissue-specific Cre PAR4KO mouse lines to investigate this very question.

Another major limitation of the study presented above is that the mice were pre-dosed with vehicle or a PAR4 antagonist before IRI. This study would need to be repeated, with the PAR4 antagonist administered post-injury, to demonstrate translational application. This study simulates prevention rather than therapeutic potential. In the clinic, doctors will prescribe a drug to treat AKI once symptoms are presented. Therefore, a study is needed to determine whether post-injury PAR4 antagonism provides the same protective benefit as seen in the previously described study. However, there is potential for an AKI drug to be administered prior to major surgery, as AKI symptoms seem to be acquired post major cardiovascular surgeries. Most clinicians are hesitant to put patients on anti-platelet/anti-coagulants before major surgeries due to the increased risk of bleeding, but PAR4 antagonism does not increase bleeding liability^35^.

Other limitations to the study presented above include the utilization of only male mice, one model of AKI by IRI, and a single strain of wild type mice. To increase the rigor of this science, both male and female mice should be studied, as well as different models of AKI, including models that simulate direct trauma to the kidney and lack of waste removal^36^.

Given the lack of an effective therapeutic for AKI and the tremendous burden of AKI on the medical system, the need for new therapeutic intervention is of great importance^1,2^. This study presents strong evidence that PAR4 is a potential therapeutic target for AKI. Not only did the VU6073819-treated mice have a protective phenotype in kidney function and inflammatory response, but there was a significant increase in survival rate compared to vehicle treated mice.

## References

1 Gameiro, J., Fonseca, J. A., Outerelo, C. & Lopes, J. A. Acute Kidney Injury: From Diagnosis to Prevention and Treatment Strategies. Journal of Clinical Medicine 9, 1704 (2020).

2 McWilliam, S. J. et al. The complex interplay between kidney injury and inflammation. Clin Kidney J 14, 780–788 (2021). 10.1093/ckj/sfaa164

3 Coughlin, S. R. Thrombin signalling and protease-activated receptors. Nature 407, 258–264 (2000). 10.1038/35025229

4 Coughlin, S. R. Protease-activated receptors in hemostasis, thrombosis and vascular biology. Journal of Thrombosis and Haemostasis 3, 1800–1814 (2005). 10.1111/j.1538-7836.2005.01377.x

5 Andrianova, I., Kowalczyk, M. & Denorme, F. Protease activated receptor-4: ready to be part of the antithrombosis spectrum. Curr Opin Hematol 31, 238–244 (2024). 10.1097/moh.0000000000000828

6 Asfaha, S. et al. Protease-activated receptor-4: a novel mechanism of inflammatory pain modulation. British journal of pharmacology 150, 176–185 (2007). 10.1038/sj.bjp.0706975

7 Bao, Y. et al. New insights into protease-activated receptor 4 signaling pathways in the pathogenesis of inflammation and neuropathic pain: a literature review. Channels (Austin) 9, 5–13 (2015). 10.4161/19336950.2014.995001

8 Gomides, L. F. et al. Blockade of proteinase-activated receptor 4 inhibits neutrophil recruitment in experimental inflammation in mice. Inflammation research : oficial journal of the European Histamine Research Society … [et al.] 63, 935–941 (2014). 10.1007/s00011-014-0767-8

9 Houle, S., Papez, M. D., Ferazzini, M., Hollenberg, M. D. & Vergnolle, N. Neutrophils and the kallikrein-kinin system in proteinase-activated receptor 4-mediated inflammation in rodents. British journal of pharmacology 146, 670–678 (2005). 10.1038/sj.bjp.0706371

10 Kahn, M. L., Nakanishi-Matsui, M., Shapiro, M. J., Ishihara, H. & Coughlin, S. R. Protease-activated receptors 1 and 4 mediate activation of human platelets by thrombin. J Clin Invest 103, 879–887 (1999). 10.1172/JCI6042

11 Erreger, K. et al. Role of protease-activated receptor 4 in mouse models of acute and chronic kidney injury. Am J Physiol Renal Physiol 326, F219–f226 (2024). 10.1152/ajprenal.00162.2023

12 Kendrick, N. C. et al. Neutrophil cathepsin G potentiates biased signaling through Protease Activated Receptor 4. Blood Advances (2025). 10.1182/bloodadvances.2025017609

13 Rosales, C. Neutrophil: A Cell with Many Roles in Inflammation or Several Cell Types? Frontiers in Physiology Volume 9 - 2018 (2018). 10.3389/fphys.2018.00113

14 Schrottmaier, W. C. & Assinger, A. The Concept of Thromboinflammation. Hamostaseologie 44, 21–30 (2024). 10.1055/a-2178-6491

15 Chen, J. et al. YAP Activation in Renal Proximal Tubule Cells Drives Diabetic Renal Interstitial Fibrogenesis. Diabetes 69, 2446–2457 (2020). 10.2337/db20-0579

16 Dabek, M. et al. Intracolonic infusion of fecal supernatants from ulcerative colitis patients triggers altered permeability and inflammation in mice: Role of cathepsin G and protease-activated receptor-4. Inflammatory Bowel Diseases 17, 1409–1414 (2010). 10.1002/ibd.21454

17 Dabek, M. et al. Luminal cathepsin g and protease-activated receptor 4: a duet involved in alterations of the colonic epithelial barrier in ulcerative colitis. Am J Pathol 175, 207–214 (2009). 10.2353/ajpath.2009.080986

18 Peach, C. J., Edgington-Mitchell, L. E., Bunnett, N. W. & Schmidt, B. L. Protease-activated receptors in health and disease. Physiological Reviews 103, 717–785 (2023). 10.1152/physrev.00044.2021

19 Dabek, M. et al. Luminal Cathepsin G and Protease-Activated Receptor 4: A Duet Involved in Alterations of the Colonic Epithelial Barrier in Ulcerative Colitis. The American Journal of Pathology 175, 207–214 (2009). 10.2353/ajpath.2009.080986

20 Fang, H. et al. Blocking protease-activated receptor 4 alleviates liver injury induced by brain death. Biochemical and Biophysical Research Communications 595, 47–53 (2022). 10.1016/j.bbrc.2022.01.074

21 Hamilton, J. R., Frauman, A. G. & Cocks, T. M. Increased Expression of Protease-Activated Receptor-2 (PAR2) and PAR4 in Human Coronary Artery by Inflammatory Stimuli Unveils Endothelium-Dependent Relaxations to PAR2 and PAR4 Agonists. Circulation Research 89, 92–98 (2001). 10.1161/hh1301.092661

22 Hollenberg, M. D., Saifeddine, M., Al-Ani, B. & Gui, Y. Proteinase-activated receptor 4 (PAR4): action of PAR4-activating peptides in vascular and gastric tissue and lack of cross-reactivity with PAR1 and PAR2. Can J Physiol Pharmacol 77, 458–464 (1999).

23 Houle, S., Papez, M. D., Ferazzini, M., Hollenberg, M. D. & Vergnolle, N. Neutrophils and the kallikrein–kinin system in proteinase-activated receptor 4-mediated inflammation in rodents. British journal of pharmacology 146, 670–678 (2005). 10.1038/sj.bjp.0706371

24 Knauss, E. A. et al. Mice with reduced protease-activated receptor 4 reactivity show decreased venous thrombosis and platelet procoagulant activity. J Thromb Haemost 23, 1278–1288 (2025). 10.1016/j.jtha.2024.12.031

25 Luo, J. et al. Antagonism of Protease-Activated Receptor 4 Protects Against Traumatic Brain Injury by Suppressing Neuroinflammation via Inhibition of Tab2/NF-κB Signaling. Neurosci Bull 37, 242–254 (2021). 10.1007/s12264-020-00601-8

26 Mao, Y., Zhang, M., Tuma, R. F. & Kunapuli, S. P. Deficiency of PAR4 attenuates cerebral ischemia/reperfusion injury in mice. J Cereb Blood Flow Metab 30, 1044–1052 (2010). 10.1038/jcbfm.2009.283

27 Rigg, R. A. et al. Protease-activated receptor 4 activity promotes platelet granule release and platelet-leukocyte interactions. Platelets 30, 126–135 (2019). 10.1080/09537104.2017.1406076

28 Sambrano, G. R. et al. Cathepsin G activates protease-activated receptor-4 in human platelets. J Biol Chem 275, 6819–6823 (2000). 10.1074/jbc.275.10.6819

29 Stoller, M. L. et al. Neutrophil cathepsin G proteolysis of protease-activated receptor 4 generates a novel, functional tethered ligand. Blood Adv 6, 2303–2308 (2022). 10.1182/bloodadvances.2021006133

30 Mao, Y., Jin, J., Daniel, J. L. & Kunapuli, S. P. Regulation of plasmin-induced protease-activated receptor 4 activation in platelets. Platelets 20, 191–198 (2009). 10.1080/09537100902803635

31 McDougall, J. J. et al. Triggering of proteinase-activated receptor 4 leads to joint pain and inflammation in mice. Arthritis & Rheumatism 60, 728–737 (2009). 10.1002/art.24300

32 Yiu, W. H. et al. Tissue kallikrein mediates pro-inflammatory pathways and activation of protease-activated receptor-4 in proximal tubular epithelial cells. PLoS One 9, e88894 (2014). 10.1371/journal.pone.0088894

33 Webb, E. M., Seymour, K. & Hamm, H. E. Trypsin Activates Human Platelets via Proteolytic Cleavage of Protease-Activated Receptor 4. Thromb Haemost (2025). 10.1055/a-2747-9083

34 Megyeri, M. et al. Complement protease MASP-1 activates human endothelial cells: PAR4 activation is a link between complement and endothelial function. Journal of immunology 183, 3409–3416 (2009). 10.4049/jimmunol.0900879

35 Wong, P. C. et al. Blockade of protease-activated receptor-4 (PAR4) provides robust antithrombotic activity with low bleeding. Sci Transl Med 9 (2017). 10.1126/scitranslmed.aaf5294

36 Bajwa, A., Pabla, N., Scindia, Y. & Gigliotti, J. in Preclinical Animal Modeling in Medicine (eds Enkhsaikhan Purevjav, Joseph F. Pierre, & Lu Lu) (IntechOpen, 2021).

